# Brain-wide electrical dynamics encode an appetitive socioemotional state

**DOI:** 10.1101/2020.07.01.181347

**Authors:** Stephen D. Mague, Austin Talbot, Cameron Blount, Lara J. Duffney, Kathryn K. Walder-Christensen, Elise Adamson, Alexandra L. Bey, Nkemdilim Ndubuizu, Gwenaёlle Thomas, Dalton N. Hughes, Saurabh Sinha, Alexandra M. Fink, Neil M. Gallagher, Rachel L. Fisher, Yong-hui Jiang, David E. Carlson, Kafui Dzirasa

## Abstract

Many cortical and subcortical regions contribute to complex social behavior; nevertheless, the network level architecture whereby the brain integrates this information to encode appetitive socioemotional behavior remains unknown. Here we measure electrical activity from eight brain regions as mice engage in a social preference assay. We then use machine learning to discover an explainable brain network that encodes the extent to which mice chose to engage another mouse. This socioemotional network is organized by theta oscillations leading from prelimbic cortex and amygdala that converge on ventral tegmental area, and network activity is synchronized with brain-wide cellular firing. The network generalizes, on a mouse-by-mouse basis, to encode socioemotional behaviors in healthy animals, but fails to encode an appetitive socioemotional state in a ‘high confidence’ genetic mouse model of autism. Thus, our findings reveal the architecture whereby the brain integrates spatially distributed activity across timescales to encode an appetitive socioemotional brain state in health and disease.

## Introduction

Social behaviors play a critical role in survival. To appropriately regulate social behavior, mammals must integrate external sensory cues with internally generated emotional brainstates. While many mechanisms whereby the brain processes external senses such as vision and audition have been elucidated, the biological processes utilized by the brain to instantiate emotional states that drive appetitive behavioral interactions remain largely unknown. This knowledge gap exists, at least in part, because emotional states have classically been inferred in preclinical models using behavioral measurements in isolation, rather than directly measuring the brain-wide activity that underlies those emotional states in behaving animals. Furthermore, collecting spatially- and temporally-resolved *in vivo* measurements of brain activity in healthy humans, who can provide self-reports of their emotional state, remains a challenge.

Multiple brain regions contribute to complex social emotional behavior. Anterior cingulate cortex activity signals empathy in humans (Morrison et al., 2004), prefrontal cortex activity regulates social hierarchy in rodents (Wang et al., 2011), and medial dorsal thalamus plays a critical role in social appetitive behavior (Ferguson and Gao, 2018). Circuit-level interactions between regions have been shown to play a role in regulating social behavior as well. Recent rodent studies have demonstrated that ventral hippocampus➔prefrontal cortex circuits mediate social memory (Phillips et al., 2019), ventral tegmental area➔nucleus accumbens circuits encode social reward (Gunaydin et al., 2014), and prefrontal cortex➔amygdala circuits are critical for social avoidance and socially aversive learning (Allsop et al., 2018; Kumar et al., 2014; Schaich Borg et al., 2017). Human electroencephalographic (EEG) studies have also described the emergence of synchronized electrical oscillations between neocortical regions at the milliseconds time scale during social perception (Fraiman et al., 2014; Rodriguez et al., 1999), and functional magnetic resonance imaging (fMRI) studies have revealed synchronized neural activity across the brain at the seconds timescale (Sokolov et al., 2018). Together, this suggests that the brain integrates neural activity across multiple brain regions and timescales to encode appetitive social brain states. Supporting this framework, a murine *in vivo* calcium imaging study identified synchronous activity across multiple cortical and limbic regions on the 100ms-seconds timescale during exposure to social novelty (Kim et al., 2016).

We have previously observed synchronous electrical oscillations on the 10ms–100ms timescale during aversive emotional states in rats and mice (Carlson et al., 2017; Hultman et al., 2018; Schaich Borg et al., 2017). We have also found that these electrical oscillations exhibit synchronous activity with millisecond-timescale cellular firing in the brain (Carlson et al., 2014; Hultman et al., 2016). Thus, we hypothesized the existence of a broad network-level mechanism involving the synchronization of electrical oscillations that integrates cellular firing across brain regions and timescales (milliseconds to seconds) to encode an appetitive social emotional brain state. Furthermore, we hypothesized that a disruption of this network architecture may play a key role in mediating social-behavioral dysfunction in neuropsychiatric disorders such as autism spectrum disorder (ASD).

To address these questions, we implanted healthy C57BL/6J (C57) mice with recording electrodes in eight brain regions including cingulate, infralimbic, and prelimbic cortex (the anatomic subdivisions of prefrontal cortex), amygdala (basolateral and central), nucleus accumbens (core and shell), medial dorsal thalamus, central hippocampus, and ventral tegmental area (VTA). We then recorded electrical oscillations and cellular firing across these regions, concurrently, as mice performed a task used to model social exploration. Using machine learning, we integrated electrical activity across these regions and across the milliseconds to seconds timescale into what we call an electrical functional connectome (*“electome”*). By analogy to the connectome, which describes the detailed anatomical connections within a brain, the *electome* describes the detailed pattern of electrical interactions across a group of brain areas. Importantly, our machine learning approach is biologically constrained such that the resultant electome model is explainable (reflects fundamental biological processes).

By analogy to gene networks, which describe collections of genes within the genome that functionally interact, *electome* networks describe the collection of brain circuits within the *electome* that together encode distinct emotional states. Learning *electome* networks that comprise the total *electome* is typically an unsupervised process, but we augmented a supervised approach to increase relevancy to complex social behavior. After confirming that an appetitive social *electome* network we discovered synchronizes cellular firing and generalizes to new animals on a mouse-by-mouse basis, we also showed that the network is dysfunctional in a genetic mouse model of ASD. Thus, our findings reveal a new mechanism whereby the brain integrates activity across space and time to encode social behavior in health and disease.

## Results

To discover the network architecture underlying a putative appetitive social brain state, we performed multisite electrical recordings while C57 strain mice were subjected to a task modeled after a classic social preference assay (Moy et al., 2007). In this behavioral assay, mice freely explore a large arena that is divided into two chambers: a small container housing a novel age- and sex-matched C3H strain mouse is situated in one chamber, and a second container holding a novel inanimate object is situated in the other (Fig. 1a). The location of the experimental mouse is tracked throughout a ten-minute exploratory period; social preference is calculated based on the relative time spent proximal to each container (Fig. 1b). Importantly, this assay can be repeated across days with new mice and new objects substituted into the containers (Fig. 1c), enabling us to collect nearly 100 minutes of electrical recordings for each of our 28 implanted mice. As expected, mice spent substantially more time interacting with a social stimulus than an object across the recording sessions (main effect of stimulus F1, 594.8= 349.1, P<0.0001 using a 3-way repeated measures ANOVA of unequal variance comparing stimulus, sex, and session; there were no other significant main effects or interactions; see also Fig. 1c).

**Figure 1:**
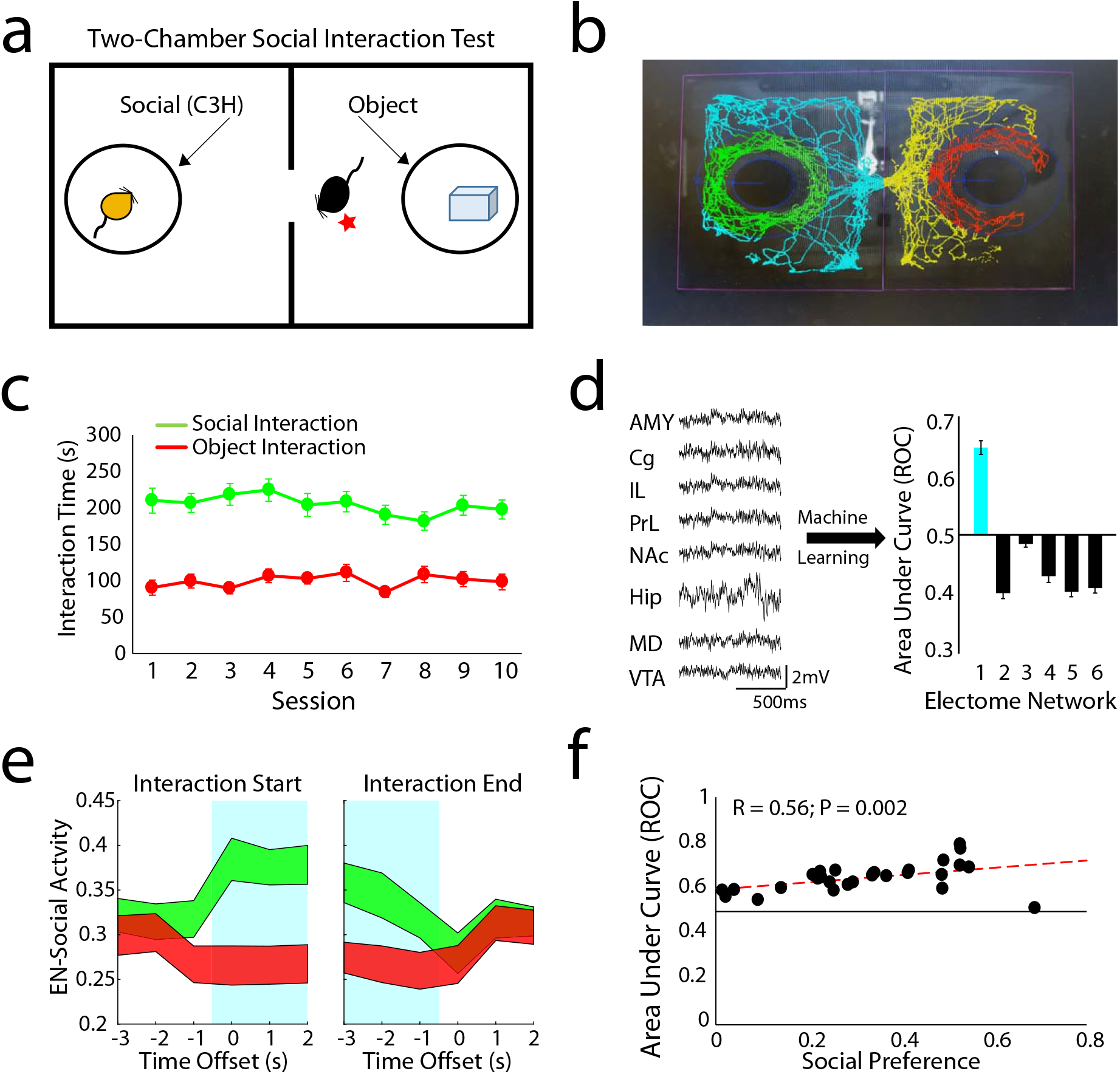
An *electome* network encodes a social emotional brain state. **a)** Schematic of the two-chamber social assay and **b)** Automated scoring approach used to quantify social and object interaction. **c)** Mice exhibit stable interaction times across repeated sessions (n=36 mice). **d)** Machine learning was used to discover six networks composed of multi-regional LFP activity [n=28 mice; amydgala (AMY), Cingulate cortex (Cg), Infralimbic cortex (IL), Prelimbic cortex (PrL), Nucleus Accumbens (NAc), Ventral Hippocampus (Hip), Medial dorsal thalamus (MD), and ventral tegmental area (VTA)]. The supervised *electome* network (blue; *EN-Social*) showed the strongest classification of social vs. object interactions. **e)** *EN-Social* event-related activity. Blue highlights identify time windows subjected to supervision by class (social vs. object). Data shown as mean±95% C.I. **f)** Decoding accuracy of *EN-Social* activity within animal versus social preference (P=0.002 using spearman correlation).

We used discriminative cross spectral factor analysis non-negative matrix factorization (dCSFA-NMF) to discover the network structure within this neural data (Talbot et al., 2020). dCSFA-NMF is a supervised machine learning approach that we designed to be both *descriptive* (i.e., discovers brain activity measures that are integrated with each other across seconds of time) and *predictive* (i.e., uses supervision to discover networked patterns of brain activity that encode external behavioral variables) (Vu et al., 2018). Importantly, dCSFA-NMF is based on widely accepted measures of brain activity, such that the resultant *electome* networks are *interpretable* (Vu et al., 2018). Specifically, each learned *electome* network integrates local field potential (LFP) power (measurement of oscillatory amplitudes across frequencies resolved from 1 to 56Hz; a neural correlate of cellular population activity and synaptic activity within brain regions), LFP synchrony (quantification of how two LFPs correlate across frequencies resolved from 1 to 56Hz over a millisecond timescale; a neural correlate of brain circuit function between brain regions), and LFP Granger synchrony (statistical forecasting based on Granger causality testing; a neural correlate of information transfer within a circuit). Finally, our dCSFA-NMF model yields an activity score for each *electome* network, which indicates the strength at which that network is represented during each one-second segment of LFP data. A given brain area or circuit can belong to multiple *electome networks*, providing the opportunity for distinct *electome networks* to functionally interact to yield a global emotional brain state (Hultman et al., 2018). Thus, dCSFA-NMF integrates spatially distributed neural activity across milliseconds to seconds of time in a manner that both models naturally occurring brain networks and predicts external behavioral conditions widely shown to induce emotional states in mice.

Using the data collected/recorded during the social preference task, we found an *electome* network that encoded social vs. object interactions. Here, we trained a model using a subset of our recorded timepoints and balanced the influence of social and object timepoints for each mouse. To encourage dCSFA-NMF to learn a network that encoded sociability (an appetitive component of social behavior), rather than solely the collection of neural responses that occurred during social interactions (e.g., sensation, arousal, etc.), we also weighted each mouse in the model based on its social preference (Supplemental Fig. S1a). Modeling our data with six *electome* networks optimally balanced complexity (i.e., explaining more variance) with parsimony (i.e., choosing fewer networks to represent the brain; see supplemental Fig. S1b). As expected, the supervised *electome* network showed the highest predictive performance (*electome* network #1, hereafter referred to as social-*electome* Network; *EN-Social*; Fig. 1d). We then probed the activity of *EN-Social* network across all the timepoints while mice explored the two-chamber assay. Though our initial learning model only used data widows labeled as social and object classes, we found that *EN-Social* activity exhibited dynamics (time course changes) that reflected behaviorally relevant task variables. Specifically, we found that ES *EN-Social* activity increased at the onset of social interactions and sloped downward as epochs of social interactions concluded. We also found that *EN-Social* activity decreased during object interactions (Fig. 1e). Thus, *EN-Social* exhibited activity that reflected multiple behavioral variables in the two-chamber social assay. Critically, the discriminatory strength of *EN-Social* was also directly correlated with social preference across our population of mice (P=0.002, R=0.56 using spearman correlation), suggesting that our weighting strategy successfully biased dCSFA-NMF to learn a network that putatively encoded an appetitive social emotional brain state.

After establishing that *EN-Social* reflects a behaviorally-relevant social brain state in the group of mice used to train our model, we explored the extent to which activity in this network mapped to widely accepted biological measures including LFP power, LFP synchrony, LFP Granger-synchrony, and cellular firing that we directly measured across our eight implanted brain regions. *EN-Social* mapped to LFP theta (4–11Hz) power within all the implanted regions. Additionally, *EN-Social* comprised prominent theta synchrony across all the implanted brain regions except ventral hippocampus (Fig. 2a, see blue highlights). The network also mapped to oscillatory activity in two higher frequency bands: 30–40Hz and 50–56Hz. The 30–40Hz oscillations showed local activity in ventral hippocampus and medial dorsal thalamus, as well as synchrony between all the implanted brain regions except ventral hippocampus (Fig. 2a, see green highlights). The higher frequency gamma band (50–56Hz) showed local activity within all the brain regions we measured except amygdala and cingulate cortex, and synchrony between all the implanted brain regions except ventral hippocampus (Fig. 2a, see red highlights). Prominent circuit directionality, quantified as the difference in the Granger synchrony between each pair of brain regions (i.e., area A➔B versus area B➔A), was observed only in the theta frequency range. This activity emerged from prelimbic cortex, infralimbic cortex, and amygdala, relayed through cingulate cortex and nucleus accumbens to medial dorsal thalamus, and converged in ventral tegmental area (see supplemental Fig. S2). Thus, *EN-Social* emerged from brain regions previously shown to play a prominent role in social behavior and converged on a brain region critical for reward regulation.

**Figure 2:**
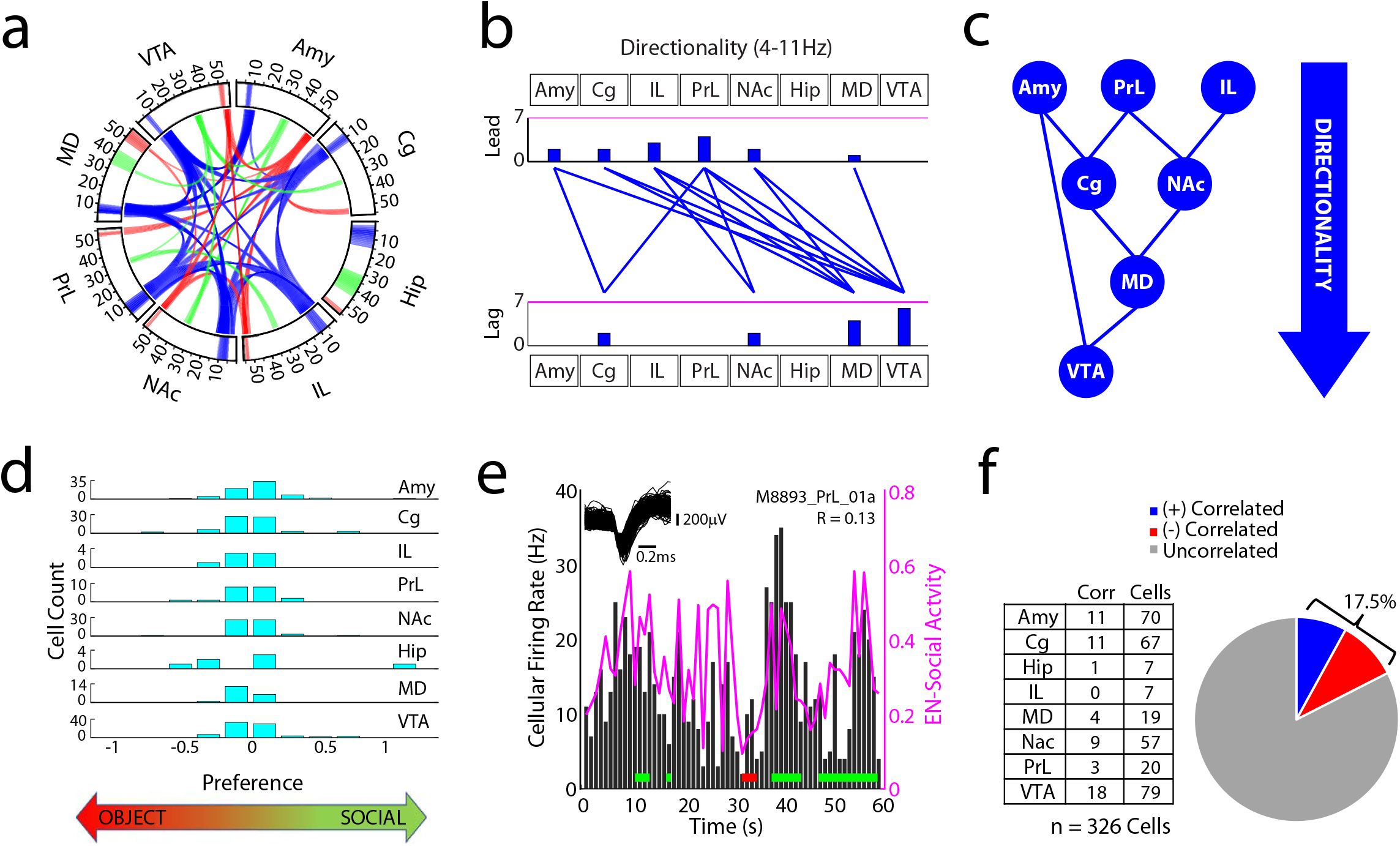
Social-emotional *electome* network maps to biological features. **a)** Power and synchrony measures that compose *EN-Social*. Brain areas and oscillatory frequency bands ranging from 1 to 56Hz are shown around the rim of the circle plot. Spectral power measures that contribute to the *electome* are depicted by the highlights around the rim, and cross spectral (i.e., synchrony) measures are depicted by the lines connecting the brain regions through the center of the circle (*electome* activity is shown at a relative spectral density threshold of 0.33, signifying the 85^th^ percentile of retained features). **b)** Granger offset measures were used to quantify directionality within the *electome* network. Prominent directionality was observed across the theta (4–11Hz) frequency band (shown at a spectral density threshold of 0.33). Histograms quantify the number of lead and lagging circuit interactions for each brain region. **c)** Schematic of signal directionality within *EN-Social*. **d)** Cellular firing preference for object vs. social interactions during two-chamber assay (cellular activity analyzed from session #5). Significant differences were observed between the two conditions for 112/326 cells (P<0.05 using rank-sum test). **e)** Representative example of cell that showed activity correlated with *EN-Social*. Horizontal red and green lines signify object and social interactions, respectively**. f)** Cellular firing vs. *EN-Social* activity across the multi-regional population of cells (P<0.05 using permutation test; recorded from session #5 of two-chamber assay).

Having identified the oscillatory signatures underlying *EN-Social*, we next verified that this *electome* network was a *bona fide* representation of biological activity and not simply an abstract mathematical construct (Hultman et al., 2018). To achieve this, we determined whether *EN-Social* activity demonstrated a relationship with the activity of cells recorded simultaneously from the implanted brain regions, an undisputed reflection of biological function. Since we found that cellular firing was broadly related to social vs. object interactions in the two-chamber assay (112/326 cells, see Fig. 2d), we used a permutation test (see methods) to rigorously test our findings. Specifically, we found that *EN-Social* exhibited a relationship to the activity of ~18% of the cells we recorded (Fig. 2e-f). Thus, we confirmed that *EN-Social* reflects a network-level neural process that emerges from cellular firing across the brain (Carlson et al., 2014; Hultman et al., 2018).

We next established *EN-Social* as a true measure of a social emotional brain state by testing the generalizability of this network, a gold-standard machine-learning validation strategy (Vu et al., 2018). Specifically, rather than simply testing whether *EN-Social* encoded object vs. social interactions in the same group of animals performing additional sessions of the two-chamber behavioral assay, we tested whether the *electome* network we learned in the initial group of mice generalized to a new cohort of C57 mice performing a different behavioral assay that also quantifies social behavior (Fig. 3a). Likewise, we also examined *EN-Social* activity in two orthogonal behavioral tasks to test whether this network was indeed encoding an emotional state relevant to the valence of external stimuli. To emphasize, *the machine learning electome model was completely blind to these tasks and data*, so this represents a test of its generalizability. First, we acquired neural activity in eight new mice exposed to our <u>F</u>ree <u>o</u>bject/<u>s</u>ocial <u>i</u>nteraction <u>t</u>est (FOSIT; Fig. 3b). In this assay, the C57 subject mice are repeatedly exposed to a novel object or a novel conspecific (age- and sex-matched) stimulus mouse. Encounters in FOSIT occur in the absence of sub chambers used for the two-chamber assay, such that the experimental mouse can also initiate social interactions with the stimulus mouse. When we projected neural data obtained during FOSIT into our initial *electome* model, we found that *EN-Social* activity was higher in reciprocal social interactions than it was in the object condition (χ^2^_3,39_ = 20.52 and P=1.3×10^-4^ using Freidman’s test; P=0.002 using post-hoc two-tailed sign-rank test with false discover rate correction; n=10 new mice). *EN-Social* activity was also higher during non-reciprocated social interactions initiated by the experimental partner mouse than it was during periods when the two mice were not interacting (P=0.01 using two-tailed sign-rank test). Finally, *EN-Social* activity tended to be higher during reciprocal interactions than the unilateral interactions, though these differences did not reach statistical significance (P=0.06 using two-tailed sign-rank test).Together, our findings verified that *EN-Social* generalized to new mice performing a different appetitive social task.

**Figure 3:**
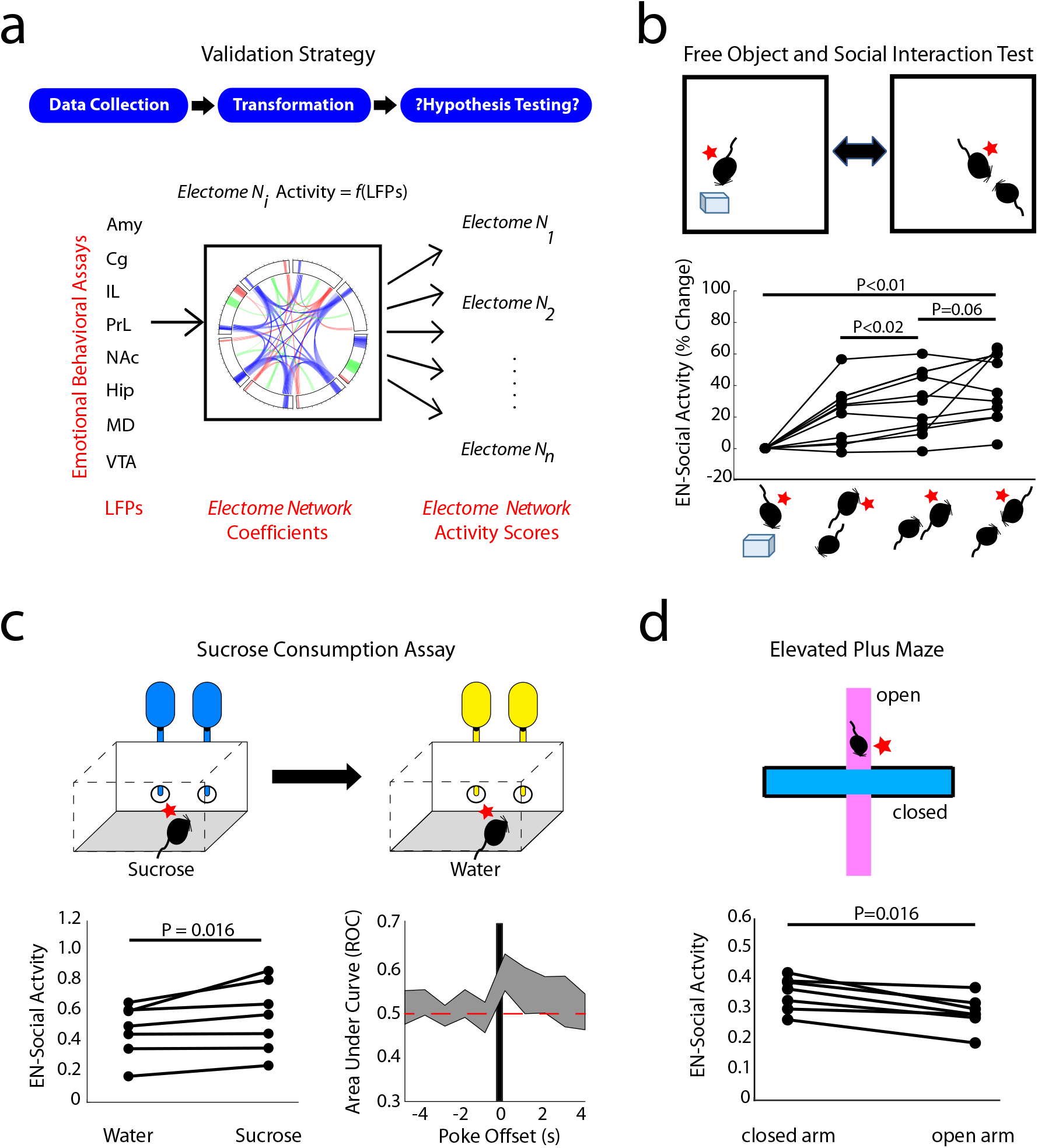
*Electome* network selectively generalizes to appetitive social-emotional brain states. **a)** Strategy for validating social *electome* network (*EN-Social*). **b)** Activity in the *EN-Social* network increased during distinct social appetitive brain states (n = 10 new mice; P<0.05 using Friedman’s test, and post-hoc testing using sign-rank test with false discover rate correction). The position of the subject mouse is shown relative to an object or another experimental mouse on the bottom. **c)** *EN-Social* activity encoded the difference between water and sucrose consumption (n=6 new mice) and **d)** encoded (negatively) the difference between the open and closed arms of an elevated plus maze (n=7 mice, 2 of which were new to the study).

We then tested whether *EN-Social* indeed encoded an appetitive state (rather than simply sensory information or salience) by testing whether the network signaled valence in other orthogonal behavioral tasks. First, we subjected a new cohort of C57 mice to an intermittent sucrose access test designed to model an appetitive state associated with food reward. Here, new mice were implanted with microwire electrode arrays and individually housed in an arena fitted with two nose poke holes. A syringe placed in the back of each hole dispensed 10uL water in response to a nose poke. After several days of habituation, water vials were replaced with 2% sucrose for 1.5 hours during the dark cycle (beginning two hours after lights off). Neural recordings were acquired during intermittent access to sucrose and the subsequent water consumption period and then projected into our initial *electome* model. *EN-Social* activity was higher following nose pokes for sucrose than for water (Fig. 3C; P=0.016 using sign-rank test; n=7 new mice), and *EN-Social* encoded the sucrose vs. water conditions to the same extent that it encoded the social vs. object conditions in the FOSIT (U=129; P=0.48 using rank-sum test; AUC = 0.59±0.02 and 0.61±0.02, for *EN-Social* in the sucrose vs. water condition and the social vs. object condition in the FOSIT. Thus, activity in *EN-social* also encoded reward in a behavioral context that was unrelated to social behavior, demonstrating that *EN-social* activity more generally reflected an appetitive brain state.

Second, we probed whether *EN-Social* activity encoded the location of mice on a classic elevated plus maze assay used to model avoidance behaviors. In this assay, mice are placed on a large plus-shaped platform that is elevated off the floor. Two of the arms of the maze are walled and the other two are open. The time that mice spend in the open arms of the maze is increased by myriad anxiolytic manipulations and decreased by anxiogenic manipulations (Krishnan et al., 2007; Marcinkiewcz et al., 2016; Rodgers et al., 1992). Furthermore, several anxiety-related neural signatures are observed as animals explore the open arms of the maze (Felix-Ortiz et al., 2016; Padilla-Coreano et al., 2016; Padilla-Coreano et al., 2019; Seidenbecher et al., 2003); thus, the open arms of this assay are widely accepted as an environmental context that induces an aversive anxiety-like brain state in C57 mice. When we projected neural data acquired from C57 mice subjected to the elevated plus maze into our initial *electome* model, we found that *EN-Social* encoded the open vs. closed arm location of mice [P=0.016 using twotailed sign-rank test; n=7 mice (5 mice from training set in a new, untrained-on behavioral condition and 2 new mice)]. Strikingly, *EN-social* activity was lower in the open arm than the closed arm (AUC=0.40±0.03 for open arm vs. closed arm, with an AUC below .5 signifying a negative relationship but the same strength as an AUC=0.60 relationship), demonstrating that network did not simply encode a brain state related an animal’s arousal state or the salience of a sensory cues. Rather, since the strength of *EN-Social* encoding was the same for the FOSIT and elevated plus maze test assays (U=73; P=0.84 for comparisons of |AUC-0.5| for the two tasks using a rank-sum test), our findings showed that EN-social encoded a brain state related to the emotional valence of external stimuli. Taken together, these results show that *EN-Social* encodes social emotional brain states in a manner that generalizes across mice and task. Furthermore, this network encodes information related to the valence state of the animal.

After demonstrating that EN-Social encoded a social appetitive brain state, we next established that the brain state was indeed causal. Here, we employed a strategy that enabled selective manipulation of neural activity within in a key node of *EN-Social* network during concurrent neurophysiological recordings and behavioral assessments. We targeted the prelimbic cortex➔nucleus accumbens circuit element, a component of *EN-social* (4-11Hz, see Fig. 2A-C), because a prior causal optogenetic had study implicated this circuit in appetitive social behavior. We implanted mice with recording electrodes and bilateral stimulating fibers in nucleus accumbens eight weeks after mice were infected with channel rhodopsin-2 (AAV5-CamKII-Chr2, Fig. 4a) in prelimbic cortex, bilaterally (n=10); thus, targeting prelimbic cortex terminals in nucleus accumbens. Four–six weeks following surgical recovery, we subjected animals to our FOSIT assay during stimulations with blue light (473nm) to activate Chr2, or yellow light (589nm) as a negative control (10Hz, 1mW bilaterally, 5ms pulse-width; Fig. 4b). Critically, we confirmed activation of the prelimbic cortex➔nucleus accumbens terminals in all of the experimental animals (Fig. 4c, left), and we excluded mice that exhibited pronounced local oscillatory responses to blue light stimulation across all of the implanted brain regions (Fig. 4c, right; n=2) given our prior observations that supraphysiological optogenetic stimulation can suppress network level activity (Hultman et al., 2018).

**Figure 4.**
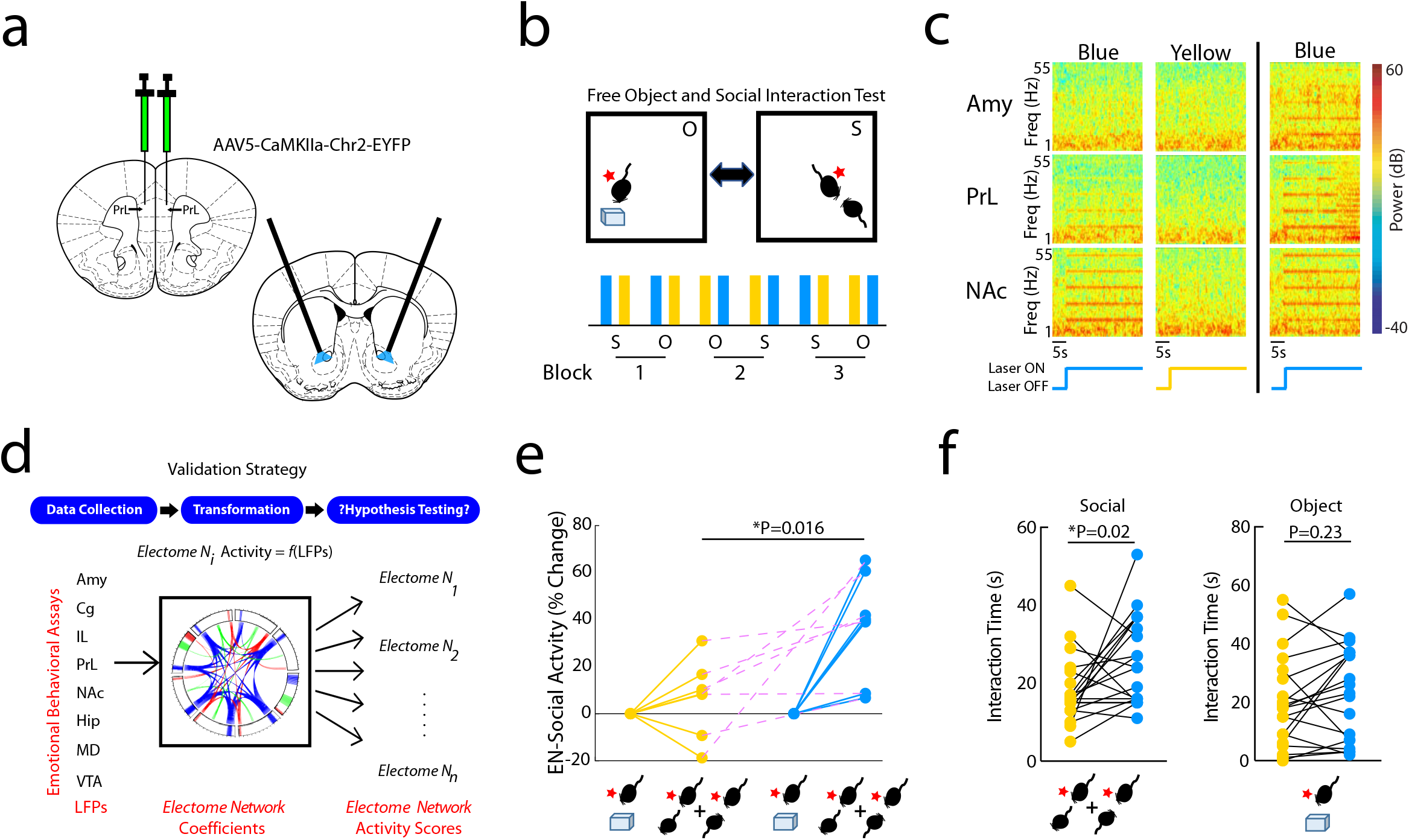
Causal activation of the prefrontal cortex to nucleus accumbens circuit element enhances *EN-Social* activity. **a)** Targeting strategy used to activate Prelimbic cortex terminals in Nucleus accumbens. **b)** Experimental paradigm for FOSIT. **c)** Power spectral plots showing increased 10Hz oscillatory activity during blue light stimulation. Plots show representative spectral patterns from a mouse during blue (left) and yellow (middle) light stimulation trials included in analysis. Representative plots from a mouse that showed increased 10Hz activity across all brain regions during blue light stimulation is shown to the right. **d)** Strategy used for *EN-Social* validation. **e)** EN-Social activity during blue light stimulation. Network activity was pooled across periods of social interaction by the subject mice and compared between the blue and yellow light stimulation periods. **f)** Social (left; P<0.05) and object interaction time (right; P>0.05) during blue and yellow light stimulation.

Causal activation of the prelimbic cortex➔nucleus accumbens circuit at 10Hz enhanced *EN-Social* activity and increased social behavior. Specifically, we projected LFP data into our initial *electome* model and quantified *EN-Social* activity during periods of social interaction (Fig. 4d). We found that blue light stimulation enhanced *EN-Social* activity compared to yellow light stimulation (P=0.016 using sign-rank test; n=7 mice; Fig. 4e). Next, we compared the amount of time mice spent socially interacting during periods of blue and yellow light stimulation. We found that blue light stimulation increased social interaction time in the FOSIT (F1,13=5.76; P=0.03 for stimulation effect using two-way RMANOVA; Fig. 4f, left). No differences in object interaction time were observed for the two light stimulation conditions (F1,13=1.67; P=0.22 for stimulation effect using two-way RMANOVA; Fig. 4f, right). Thus, our findings showed that activation of the prelimbic cortex➔nucleus accumbens circuit enhanced both *EN-Social* activity and increased social interaction. Taken together with our other validation experiments, these results provided strong evidence that *EN-Social* was causally related to appetitive social behavior.

After establishing *EN-Social* as a generalized and putatively causal appetitive social emotional brain state under healthy conditions, we wondered whether the appetitive aspects of *EN-Social* would be altered in a disease state associated with social deficits. Autism spectrum disorder (ASD) is a pervasive neurodevelopmental disorder for which social deficits are a core feature. They include deficits in social attention and engagement and deficient processing of social information (Crawford et al., 2016; Dawson et al., 2012; Dawson et al., 2004; Klin et al., 2015). Genetic manipulations are implicated in ~52% of ASD cases (Gaugler et al., 2014), and one such high confidence gene is ANK2 which codes the Ankyrin-B protein (SFARI-GENE, 2020; Yang et al., 2019). Importantly, unlike many other genes that are implicated in syndromic ASD, ANK2 mutations yield social deficits without impacting executive cognitive dysfunction. We previously developed an ANK2 mouse model based on a gene mutation initially identified in a patient with ASD showing social deficits with normal IQ and no seizures. Heterozygous mice show decreased social behavior on multiple assays, decreased juvenile vocalizations, and increased cognitive flexibility (Yang et al., 2019).

We implanted adult ANK2 male mice and their wild-type littermate controls with recording electrodes and subjected them to neural recordings in the two-chamber social assay (Fig. 5a-b). ANK2 mice exhibited normal social preference (U=64; P=0.86 using rank-sum test; Fig. 5c) and did not show seizure activity (Fig. 5d-e). When we projected their LFP activity into our initial *electome* model, ANK2 mice and their littermate controls both exhibited *EN-Social* activity that was higher during social vs. object encounters (F1,16=30.5; P=4.7×10^-5 for social vs. object effect using a mixed-model ANOVA; n=11 and 7, for wild-type and ANK2 mice, respectively; Fig. 5f). Furthermore, no differences in *EN-Social* activity were observed across genotype (F1,6=0.58; P=0.46 for genotype effect; F1,16=1.04; P=0.32 for interaction effect), demonstrating that *EN-Social* continued to encode socially relevant information in the mutants. We then tested whether mutants also encoded the appetitive component of *EN-Social*.

**Figure 5:**
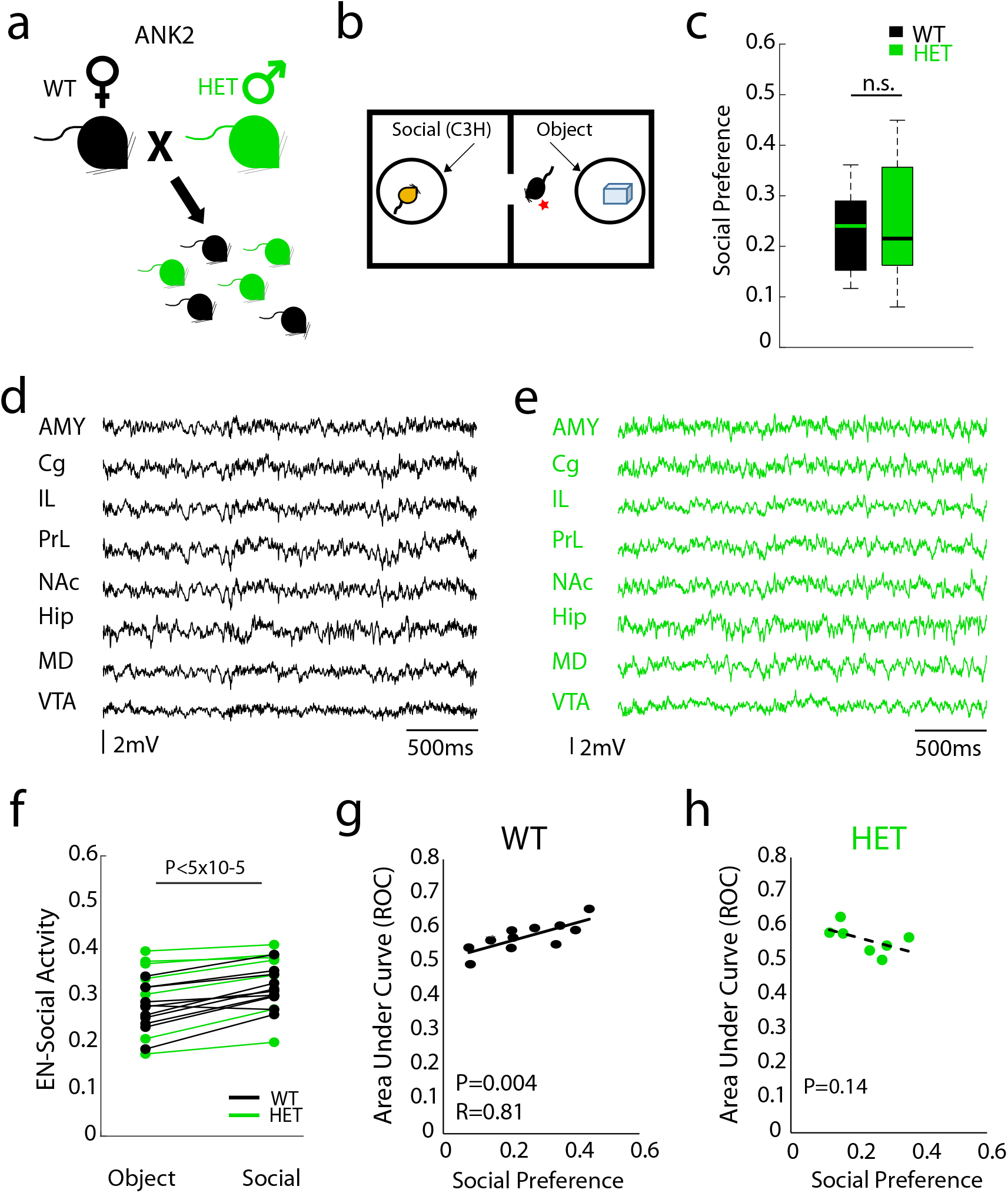
*Electome* network fails to encode social preference in a genetic model of Autism spectrum disorder. **a-b)** ANK2 mice and their littermate controls were subjected to the two-chamber social assay. **c)** Both ANK2 mice and their littermate controls showed preference for social interactions (P>0.05). **d-e)** Representative LFP activity in d) wild type and e) ANK2 mice showing no seizure activity. **f)** EN-Social activity during social and object interactions (P<0.05 for conditions; P>0.05 for genotype effects). **g-h)** *EN-Social* activity vs. appetitive social behavior in g) wild-type mice (P<0.05) and h) ANK2 mutants (P>0.05).

The discriminatory strength of *EN-Social* was directly correlated with social preference on a mouse-by-mouse basis across the group of wild-type mice (F_1,14_=10.1; P=0.007 for interaction effect using Analysis of Covariance; P=0.004 and RHO=0.81 for wild-type mice using spearman rank test; see Fig. 5g), demonstrating that the socially appetitive brain state we discovered in our original group of training animals generalized to new mice. Strikingly, *EN-Social* activity was not correlated with social preference in the ANK2 mutants (P=0.14 for ANK2 mutants using spearman rank test; see Fig. 5h). Thus, *EN-Social* failed to encode the socially appetitive brain state in ANK2 mice, confirming that *EN-Social* was altered in a disease state associated with social deficits.

## Discussion

The manner whereby cells, segregated across multiple brain regions, integrate their activity over time to generate socially appetitive brain states remains an unaddressed question. Human studies have sought to discover this network-level mechanism by probing changes in brain-wide hemodynamic responses using fMRI and/or fast electrical activity across the scalp using EEG. These studies have revealed multiple brain regions and several fast-neural oscillatory features that putatively contribute to appetitive social processing (Fraiman et al., 2014; Rodriguez et al., 1999; Sokolov et al., 2018). Nevertheless, fMRI is limited in its ability to resolve neural activity at the timescale of cellular activity in the brain (i.e. milliseconds), EEG does not quantify neural activity deep within the brain, and causality testing via direct manipulation of the human brain remains a challenge. Preclinical animal studies, on the other hand, readily facilitate causality testing of genetic and cellular/molecular mechanisms; however, approaches that monitor electrical activity across multiple regions have yet to be broadly applied to the study of emotional behavior. Given these limitations, network models that describe the causal mechanism whereby fast neural activity throughout the depth of the brain integrates across space and time to encode social-appetitive behavior remain elusive.

Here we implanted recording wires into eight cortical and limbic brain regions located through the depth of the brain, allowing us to record millisecond-timescale electrical fluctuations as mice engaged in behaviors used to model appetitive and aversive social brain states. Our neural recordings yielded 5152 features that quantified fast timescale (i.e., milliseconds to hundreds of milliseconds) region-specific activity and between-region circuit activity each second. Using machine learning, we discovered the biophysiological patterns whereby these features integrated across seconds of time to encode an appetitive social-emotional brain state. Not only did we discover that activity in the resulting *electome* network encoded the onset and termination of social interaction epochs (Fig. 1e), we also confirmed that the activation strength of *EN-Social* was correlated with the social preference of individual mice (Fig. 1f). Both these properties generalized to new groups of mice that were not used to discover the initial network on a mouse-by-mouse basis (see Fig. 4g and supplemental Fig. S3), and the network generalized across sex (unpublished findings). Strikingly, we found that *EN-Social* generalized to encode active and passive social engagement in a different task that allows two freely behaving mice to interact with each other (i.e., FOST), confirming its validity. *EN-Social* also encoded food reward. Interestingly, the network exhibited some spectral overlap (based on brain regions, frequencies, and directionality composition) with another *electome* network we recently found to signal goal progress (Vu et al., 2019), suggesting that the *EN-Social* may exploit general brain circuits which encode reward. Finally, *EN-Social* encoded an emotional state related to anxiety avoidance, demonstrating that the network signaled the valence of external stimuli.

Activity in the *electome* network correlated with cellular firing throughout the brain, confirming its biological significance. The network was composed of theta oscillations (4-11Hz) that synchronized across most of the regions we measured, and gamma oscillations that were most prominent in the 30–40Hz and 50–56Hz bands. Directionality in the network was largely organized within the theta oscillatory frequency, and this activity emerged from amygdala, prelimbic cortex, and infralimbic cortex, relayed through cingulate cortex and nucleus accumbens to medial dorsal thalamus, and converged in VTA.

Critically, directionality observed within a circuit by no means implies that information flow is unidirectional (Granger A➔B exceeding Granger B➔A does not denote Granger A➔B but not Granger B➔A), nor does directional Granger coherence preclude other regions serving as anatomic relays (Granger A➔B does not exclude Granger A➔Z➔B). Nevertheless, it is notable that the activity pattern we discovered in *EN-social* mirrored findings from other causal studies aimed at dissecting the individual anatomical circuits that contribute to social behavior. For example, while hyperactivation of prelimbic cortex projection neurons disrupts social preference (Yizhar et al., 2011), projection specific studies revealed that the prelimbic cortex ➔ nucleus accumbens circuit, but not the prelimbic cortex➔amygdala or prelimbic cortex➔VTA circuits mediates this effect (Murugan et al., 2017). This aligns with the directionality in *EN-Social* which exhibits activity in the prelimbic cortex ➔ nucleus accumbens circuit, but not the prelimbic cortex➔amygdala or prelimbic cortex➔VTA circuits.

In contrast to these previous findings, here we found that stimulation of the prelimbic cortex ➔ nucleus accumbens circuit induced social behavior. This discrepancy may be due to difference in stimulation frequency used in the prior work (10Hz and 20Hz), the context in which social encounters occurred (novel area vs. habituated arena), or difference in experimental design (within-subjects vs. cross-subjects) (Murugan et al., 2017). Nevertheless, since we found that 10Hz stimulation of this pathway also enhanced EN-Social activity, we believe that our data support a causal role of prelimbic cortex ➔ nucleus accumbens and EN-Social in supporting social behavior. Finally, prior work has also implicated the cingulate cortex➔amygdala circuit in mediating aversive social emotional states (Allsop et al., 2018). While our first *electome* model learned in a socially aversive context identified this same circuit pathway (Schaich Borg et al., 2017), the cingulate cortex➔amygdala was not prominently featured in our current appetitive social *electome* network. Thus, the network we discovered clarifies how distinct circuits integrate in a normal physiological context to encode an appetitive social emotional brain state. Notably, we found that *EN-Social* also encodes emotional valence in other orthogonal behavior tasks including sucrose drinking and the elevated plus maze. Future analysis using adversarial machine learning models may clarify whether and/or disambiguate which specific aspects of *EN-Social* uniquely signal appetitive social behavior rather than generally signal all appetitive brain states.

Our findings also established a role for altered *EN-Social* function in pathological emotional brain states. ASD is a neurodevelopmental disorder characterized by social deficits, including aberrant social attention and engagement and deficient processing of social information (Crawford et al., 2016; Dawson et al., 2012; Dawson et al., 2004; Klin et al., 2015). There is no clear genetic basis for the disorder in many individuals, and a convergent cellular/molecular pathway that globally mediates social deficits in ASD remains elusive. Conversely, network-level deficits in processing faces, facial affect, and biological motion manifest in ASD, even in this absence of known genetic underpinnings. For example, high-risk infants who later develop ASD show atypical neural responses when viewing face stimuli and dynamic complex social stimuli (Jones et al., 2016), and EEG studies demonstrate impaired face and facial affect processing in children and adults with ASD (Dawson et al., 2005; Jokisch et al., 2005; Kroger et al., 2014; Pavlova et al., 2004; Ulloa and Pineda, 2007). Together, these observations suggest that a convergent network-level mechanism, rather than a convergent cellular/molecular-level mechanism, may mediate social deficits in the disorder. Indeed, aberrant neural responses to faces and facial affect are considered amongst the most promising ASD biomarker candidates (Jeste and Geschwind, 2016; Loth et al., 2016; McPartland, 2016; Ruggeri et al., 2014).

Here, we subjected an ANK2-based mouse model of ASD to *in vivo* neural recordings during a classic social behavioral assay. The ANK2 mutants exhibited altered function of *EN-Social*. Specifically, *EN-Social* encoded the difference between social and object interactions, but it failed to encode the social preference of individual ANK2 mutant animals. Thus, though *EN-Social* successfully encoded a social brain state in the ‘high-confidence’ ASD model, the circuit components of *EN-Social* that encode socially appetitive information in healthy animals were altered in the mutants. These findings raise the intriguing potential that ANK2 mice may fail to encode an appetitive brain state, or that a different set of brain circuits or networks may sub-optimally encode socially appetitive behavior in ASD. Future experiments in which *electome* networks are trained across larger groups of ANK2 mutants may clarify this question.

ASD behavior is typically quantified in mice using a battery of social behavioral tasks. Nevertheless, it is well known that several mouse lines which exhibit genetic construct validity with the human disorder fail to exhibit measurable deficits in specific assays. For example, we found that our ANK2 mice exhibited normal social preference in our two-chamber social preference assay. This is consistent with our prior observations in this mouse line (Yang et al., 2019), and another mouse line engineered to exhibit construct validity with another high confidence risk gene, SHANK3 (SFARI-GENE, 2020; Wang et al., 2016). On the other hand, both lines exhibit clear deficits in separate social-behavioral assays, suggesting that our approach of quantifying circuit/network activity may have higher sensitivity for detecting the pathophysiological processes underlying social deficits in ASD than behavioral measurements in isolation.

Supporting this notion, our prior experiments in normal mice exposed to social stimuli identified circuit components that shared spectral features with *EN-Social* (including synchronized theta oscillations between prelimbic cortex, nucleus accumbens, cingulate cortex, and medial dorsal thalamus), and SHANK3 mutant mice showed diminished activation of several of these circuits (Wang et al., 2016). In our other work, we also found *EN-Social* dysfunction in a mouse model of ASD that was based on stress-induced manipulations of the prenatal environment (unpublished findings). These observations suggest that many causal ASD manipulations may globally disrupt the encoding of socially appetitive brain states. Alternatively, different and likely less efficient networks may emerge to encode socially appetitive brain states at key developmental timepoints. Both outcomes suggest that *EN-Social* dysfunction may reflect a convergent mechanism underlying social deficits in ASD, and additional experiments are warranted to directly test these hypotheses.

There are no pharmacological agents that treat social deficits in ASD. The identification and validation of a socially appetitive brain state in normal mice and ASD models raises the potential that brain spatiotemporal dynamics can be exploited to develop ASD diagnostics and stimulation-based therapeutics that ameliorate social dysfunction in individuals with ASD.

## Acknowledgements

We would like to thank Vann Bennett for contributing the ANK2 mutant mice; Staci Bilbo, Cagla Eroglu, Carina Block, and Rainbo Hultman for comments on this work; Derek Southwell, and Timothy Nyangacha for technical support. This work was supported by WM Keck Foundation grant to KD and Fan Wang; NIH grant R01MH120158 to KD; NIH grant R21MH104316 to KD and YHJ; NIH grant R01ES025549 to KD, SB, and CE; and NIH grant 1R01EB026937 to DEC and KD. A special thanks to Freeman Hrabowski, Robert and Jane Meyerhoff, and the Meyerhoff Scholarship Program.

