## Supplemental Materials for "Brain-wide electrical dynamics encode an appetitive socioemotional state"

### Supplemental Figures

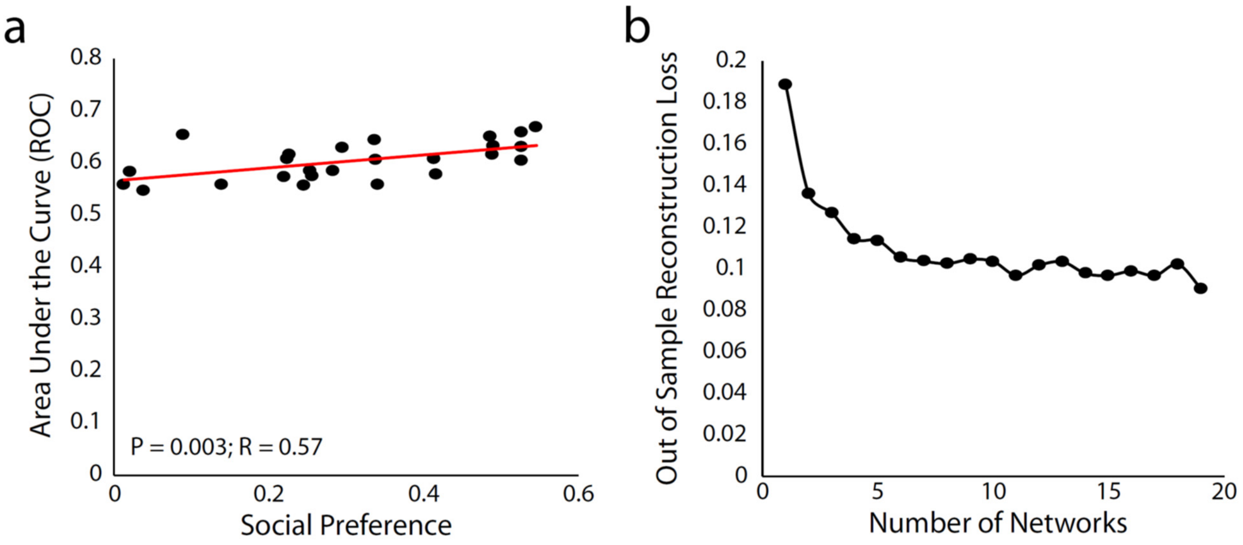


**Supplemental Figure S1: Selection of model features, related to Fig. 1. a)** We initially trained an unsupervised PCA model using power and synchrony measures across the implanted areas for each animal. We used 70% of the social and object timepoints for model training, and the remaining 30% of the data was used for hold-out testing. Since the area under the curve of the receiver operating characteristic for each animal was correlated with their social preference, we weighed individual mice within our subsequent dCSFA-NMF Electome model based on this behavioral measure. Thus, in addition to encoding the difference between social and object interactions, this approach also encouraged dCSFA-NMF to learn a network that also jointly encoded social preference. **b)** We calculated the out-of-subject reconstruction loss using the initial dCSFA-NMF model vs. the numbers of trained networks. Gains in the reconstruction loss diminished with >6 networks.

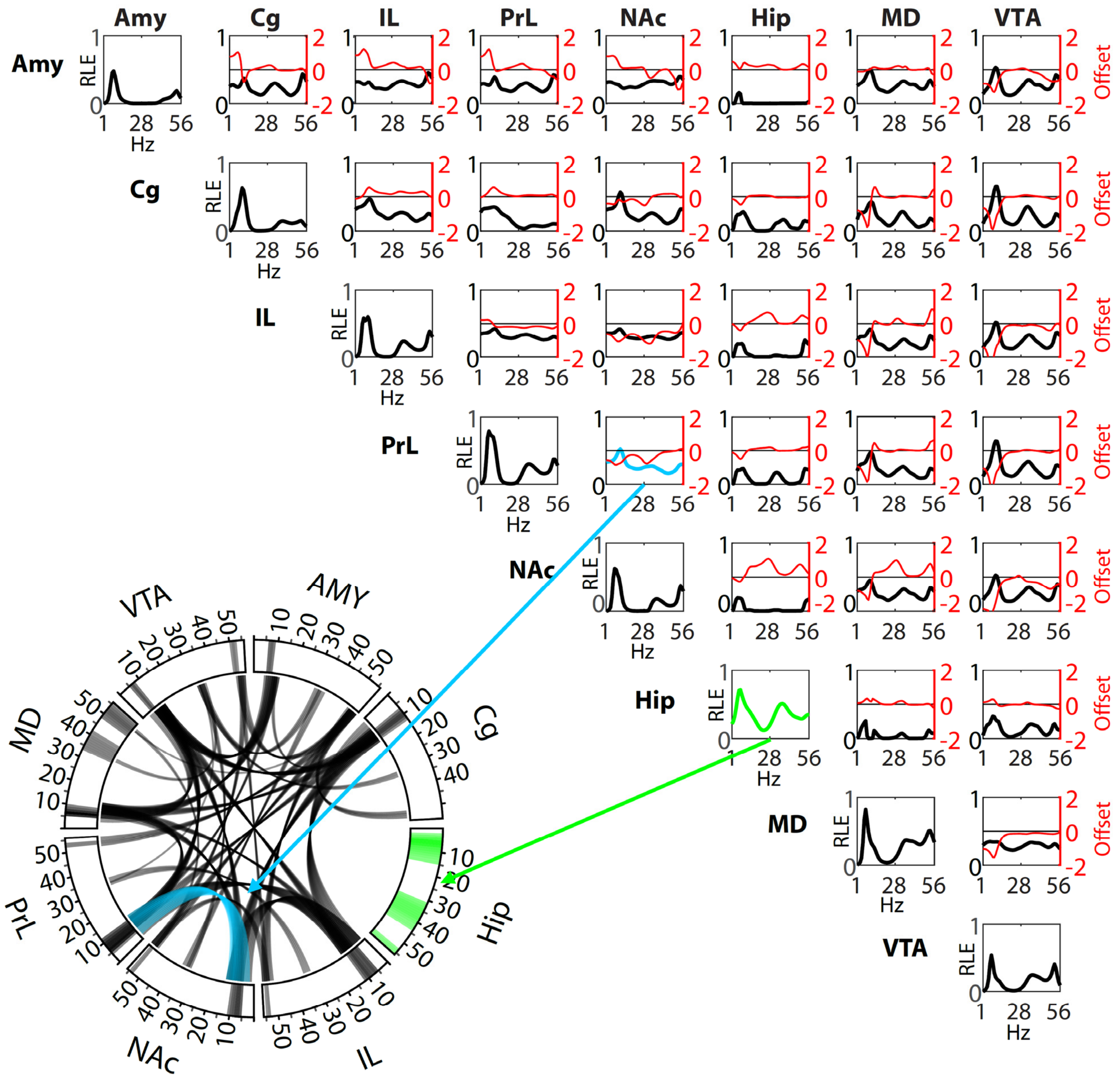


**Supplemental Figure S2: Power and synchrony measures that define the Social-*Electome* Network*,* related to Fig. 2.** Brain areas are shown to the top and the left identifying power and synchrony density functions for *Electome Factor* 1. Amplitude values (shown in black) reflect the relative LFP spectral energy (RLE) observed at each frequency, where the *Electome Factor* is normalized to the total energy observed across the 6 networks. The offset between the two non-normalized granger synchrony functions for each brain area pair (A→B and B→A) are also shown in red (i.e. directionality; axis scale to the right). Positive spectral offsets correspond to frequencies at which the area listed along the top leads the area listed on the left. Negative spectral offsets correspond to the frequencies at which the area listed on the left leads the area listed on the top. The circular plot depicts the frequencies for power (outer rim) and synchrony (curved lines connecting two regions) above an amplitude threshold of 0.33. As a representative example, the power measures for ventral hippocampus (Hip) are highlighted in bright green in both the circular and correlation plots; synchrony between prelimbic cortex (PrL) and nucleus accumbens (NAc) are highlighted in cyan.

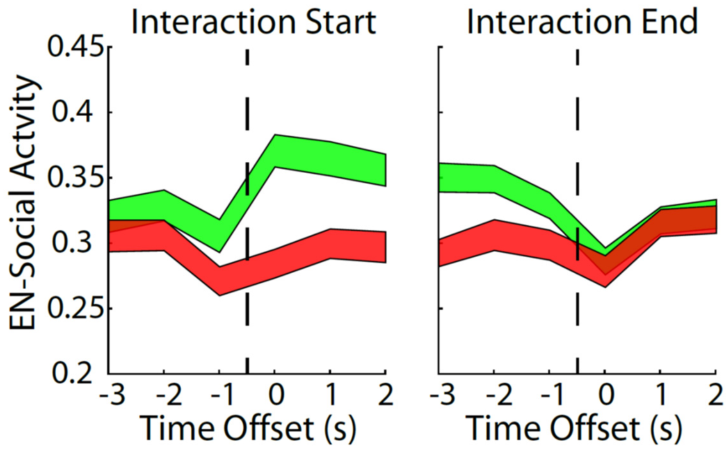


**Supplemental Figure S3: Event-related activity of the Social-*Electome* network, related to Fig. 1 and 3.** As part of our strategy outlined in Fig. 3a, twenty-eight new healthy mice were implanted with recording electrodes, and neural data was acquired during the social preference test (10 sessions). We then projected LFP data into the Social-*Electome* network model learned from the initial training animals. Note that the same behaviorally-relevant network dynamics are observed in these hold-out mice (compare to Fig. 1e). Data shown as mean±s.e.m.

**Supplemental Table S1: Extended Author Contributions**

| | |
|---|---|
| **Stephen D. Mague** | Jointly conceived two-chamber social interaction experiment; jointly built electrodes; jointly performed electrode implantations in C57BL/6J mice for two-chamber social interaction experiment with KD; jointly performed electrode implantations for sucrose drinking and elevated plus maze experiments with CB; jointly analyzed behavioral data for two-chamber social interaction experiments and performed neurophysiological data processing for all experiments presented in the paper with CB, EA, NN, and KKW; jointly conceived FOSIT experiment with KD and AT; jointly conceived and performed optogenetic stimulation experiments with KD and KKW; jointly performed histological confirmations for all experiments with LJD, GET, EA, NN CB, and KKW; jointly supervised all data collection and wrote the paper with DEC and KD. |
| **Austin Talbot** | Developed and implemented the CSFA-NMF machine learning analyses utilized for all neurophysiological analysis in the paper including the model discovery and projections of new neurophysiological data into the initial model space; edited the paper. |
| **Cameron Blount** | Built electrodes for C57BL/6J mice two-chamber social interaction test, FOSIT, EPM and ANK2 experiments. Jointly implanted mice for EPM experiment; jointly collected data for C57BL/6J mice two-chamber social interaction test experiment with LJD and NN; Collected data for FOSIT in C57BL/6J mice; jointly collected data for ANK2 experiments with KKW and ALB; jointly analyzed behavioral data for two-chamber social interaction experiments and performed neurophysiological data processing for all experiments presented in the paper with SDM, EA, NN, and KKW; jointly performed histological confirmations for all experiments with LJD, SDM, GET, EA, NN, and KKW. |
| **Lara J. Duffney** | Jointly collected data from C57BL/6J mice for two-chamber social preference experiment with CB and NN; jointly performed histological confirmations for all experiments with EA, GET, NN, CB, and KKW; jointly conceived of two-chamber social preference experiment for C57BL/6J mice with KD. |
| **Kathryn K. Walder-Christensen** | Jointly collected data for ANK2 experiments with CB and ALB; jointly conceived and performed optogenetic stimulation experiments with KD and SDM; jointly performed histological confirmations for all experiments with LJD, SDM, GET, EA, NN, and CB; edited the paper. |

| | |
|---|---|
| **Elise Adamson** | *Jointly performed behavioral and neurophysiological data processing for two chamber experiments in* C57BL/6J *with* SDM, *CB,* NN, and KKW; Conceived, collected data, and performed neurophysiological data processing for sucrose consumption experiment; jointly performed histological confirmations for all experiments with LJD, SDM, GET, NN, CB, and KKW; edited the paper. |
| **Alexandra L. Bey** | Performed all behavioral analysis for FOSIT experiment; jointly collected data for ANK2 experiments with KKW and CB; edited the paper. |
| **Nkemdilim Ndubuizu** | Jointly collected data from C57BL/6J mice for two-chamber social preference experiment with CB and LJD; jointly analyzed behavioral data for two-chamber social interaction experiments and performed neurophysiological data processing for all experiments presented in the paper with CB, EA, SDM, and KKW; jointly performed histological confirmations for all experiments with LJD, SDM, GET, EA, CB, and KKW. |
| **Gwenaëlle Thomas** | Jointly performed histological confirmations for all experiments with SDM, LJD, EA, NN CB, and KKW; edited the paper. |
| **Dalton Hughes** | Conceived, collected data, and performed neurophysiological data processing for elevated plus maze experiment. |
| **Saurabh Sinha** | Analyzed LFP data from ANK2 mice for seizure activity. |
| **Alexandra M. Fink** | Contributed to preprocessing of data for Spike-*Electome Factor* analysis. |
| **Neil M. Gallagher** | Implemented granger features for data processing; jointly contributed to data preprocessing for elevated plus maze and sucrose consumption tasks. |
| **Rachel L. Fisher** | Jointly built electrodes for ANK2 experiments; Built all optoelectrodes for optogenetics studies; Contributed to histological confirmations ANK2 experiments. |
| **Yong-hui Jiang** | Jointly supervised LD with KD. |
| **David E. Carlson** | Supervised development of all aspects of CSFA-NMF machine learning analyses and data processing methodology; wrote the paper with SDM and KD. |
| **Kafui Dzirasa** | Jointly conceived of experiments with LJD, AT, KKW, DH, EA, SDM, DEC; performed surgical implantations for all experiments presented in this paper; jointly performed optogenetic experiments with SDM and KKW; jointly analyzed neurophysiological data with AT; supervised all |

| | |
|---|---|
| | behavioral and neurophysiological experiments with SDM; wrote the paper with SDM and DEC. |

### Supplemental Methods

#### CONTACT FOR REAGENT AND RESOURCE SHARING

#### EXPERIMENTAL MODEL AND SUBJECT DETAILS

**Animal Care and Use**

C57BL/6J (C57) mice purchased from the Jackson Labs were used for two-chamber experiments used to train the initial Electome model, the subsequent studies using the free object social interaction test, the forced interaction test, the elevated plus maze, the sucrose consumption task, and the optogenetic manipulation studies. ANK2 mutant mice were generated as previously described (Yang et al., 2019). These mutants and their WT littermate controls were bred within the Duke Vivarium. C3H strain mice used for the two-chamber social interaction test were purchased from Jackson Labs. All mice were housed 3-5 per cage, maintained on a 12-hour light/dark cycle, in a humidity- and temperature-controlled room with water and food available *ad libitum*.

Studies were conducted with approved protocols from the Duke University Institutional Animal Care and Use Committee and were in accordance with the NIH guidelines for the Care and Use of Laboratory Animals. Studies were conducted using mice that were 12-20 weeks old.

#### METHOD DETAILS

**Electrode implantation surgery**

Mice were anesthetized with 1% isoflurane, placed in a stereotaxic device, and metal ground screws were secured above the cerebellum and anterior cranium. The recording bundles designed to target basolateral and central amygdala (AMY), medial dorsal thalamus (MD), nucleus accumbens core and shell (NAc), VTA, medial prefrontal cortex (mPFC), and VHip were centered based on stereotaxic coordinates measured from bregma (Amy: -1.4mm AP, 2.9 mm ML, -3.85 mm DV from the dura; MD: -1.58mm AP, 0.3 mm ML, -2.88 mm DV from the dura; VTA: -3.5mm AP, ±0.25 mm ML, -4.25 mm DV from the dura; VHip: -3.3mm AP, 3.0mm ML, -3.75mm DV from the dura; mPFC: 1.62mm AP, ±0.25mm ML, 2.25mm DV from the dura; NAc: 1.3mm AP, 2.25mm ML, -4.1 mm DV from the dura, implanted at an angle of 22.1°). We targeted cingulate cortex, prelimbic cortex, infralimbic cortex using the mPFC bundle by building a 0.5mm and 1.1mm DV stagger into our electrode bundle microwires. Animals were implanted bilaterally in mPFC and VTA. All other bundles were implanted in the left hemisphere. The NAc bundle included a 0.6mm DV stagger such that wires were distributed across NAc core and shell. We targeted BLA and CeA by building a 0.5mm ML stagger and 0.3mm DV stagger into our AMY electrode bundle. In order to mitigate pain and inflammation related to the procedure, all animals received carprofen (5 mg/kg, s.c.) injections once prior to surgery and then once every 24 hours for three days following electrode implantation.

**Histological Confirmation**

Histological analysis of implantation sites was performed at the conclusion of experiments to confirm recording sites used for neurophysiological analysis. Animals were perfused with 4% paraformaldehyde and brains were harvested and stored for 24 hrs in PFA. Brains were cryoprotected with sucrose and frozen in OCT compound and stored at -80C. Brains were sliced at 35µm and stained using either DAPI (ab104139, AbCam, Cambridge, MA), NeuroTrace fluorescent Nissl Stain (N21480, ThermoFisher Scientific, Waltham, MA) or cresyl violet (C5042, Sigma-Aldrich, St. Louis, MO) using standard protocols. Images were obtained using a Nikon Eclipse fluorescence microscope at 4x and 10x magnifications. We took the following approaches to histological confirmation. When we performed complete histological analysis on 54 animals, we found 11/432 mistargeted implants (2.5% error rate). We observed a similar error rate (~3%) after complete histological analysis on an additional 56 mice. Since machine learning analysis benefits from larger data sets and can be more robust to data variance than classic frequentist statistics, we employed the following strategy. To learn our *EN*-Social model, we concluded that a training set containing 27/28 accurate data points per region was more desirable than a training set that contained 21/21 accurate data points per region. Thus, we used all 28 implanted animals to learn our initial model. We employed a similar strategy for our validation analysis. Specifically, presuming accurate targeting with 97% certainty, we included animals with missing or damaged histological slices in our analysis. However, if there was clear histological confirmation of mistargeting for any of the recorded regions, the animal was removed.

**Neurophysiological data acquisition**

Mice were connected to a headstage (Blackrock Microsystems, UT, USA) without anesthesia, and placed in each behavioral arena. Neuronal activity was sampled at 30kHz using the Cerebus acquisition system (Blackrock Microsystems Inc., UT). Local field potentials (LFPs) were bandpass filtered at 0.5–250Hz and stored at 1000Hz. An online noise cancellation algorithm was applied to reduce 60Hz artifact. Neuronal data were referenced online against a wire within the same brain area that did not exhibit a SNR > 3:1. At the end of the recording, cells were sorted again using an offline sorting algorithm (Plexon Inc., TX) to confirm the quality of the recorded cells. Only cellular clusters well-isolated with respect to background noise, defined as a Mahanalobis distance greater than 3 compared to the null point, were used for our unit-Electome Factor correlation analysis. Clusters that exhibited more than 99% of their inter-spike-interval distribution above 2ms were defined as single units (93.5% of recorded neurons). Ultimately, we chose to use both single and multi-units for our analysis since our sole objective was to determine whether the Electome Network activity showed temporal dynamics that reflected cellular activity. This strategy mirrors our prior experiments probing the network level mechanisms underlying depression vulnerability (Hultman et al., 2018). Neurophysiological recordings were referenced to a ground wire connected to both ground screws.

**LFP preprocessing to remove signal artifact**

We used a heuristic to remove recording segments with non-physiological signals. First, we estimated the envelope of the signal in each channel using the magnitude of the Hilbert transform. For any 1-second window where the envelope exceeds above a pre-selected low threshold, the entire segment is removed if the envelope exceeds a second, high threshold at any point within that window. The two thresholds were determined independently for each brain region. The high threshold was selected to be 5 times the median absolute deviation of the envelope value for that region. Five median absolute deviations was chosen as the high threshold because it is roughly equivalent to 3 standard deviations from the mean for normally distributed data, but is robust to outliers in the data. The low threshold was empirically chosen to be 3.33% of the high threshold. If more than half the window was removed for a channel, we removed the rest of that window for that channel as well. In addition, any windows where the standard deviation of the channel is less than 0.01 were also removed. Using this approach, 13±3.5% of the data/mouse (N=28 for our model training) were excluded from this analysis. This conservative strategy optimized the potential of our learning model to discover a network that was uniquely related to appetitive social emotional brain states.

**Determination of LFP oscillatory power and cross-area synchrony and granger coherence**

LFPs were averaged across wires within region to yield a composite LFP measure. Signal processing was performed using Matlab (The MathWorks, Inc., Natick, MA). For LFP power, a sliding Fourier transform with Hamming window was applied to the averaged LFP signal using a 1 second window and a 1 second step. Frequencies were analyzed with a resolution of 1Hz. LFP cross-structural coherence was calculated from the pairs of averaged LFPs using magnitude-squared coherence

$$C_{AB}(f) = \frac{|Psd_{AB}(f)|^2}{Psd_{AA}(f)Psd_{BB}(f)}$$

where coherence is a function of the power spectral densities of A and B, and their cross-spectral densities.

The spectral Granger causality (Geweke, 1982) features were calculated using the *Multivariate Granger Causality* (MVGC) MATLAB toolbox (Barnett and Seth, 2014). The non-stationary data required a highpass, so a highpass Butterworth filter with a stopband at 1Hz and a passband starting at 4Hz was applied to the data. Granger causality values for each window were calculated using a 20-order AR model via the *GCCA_tsdata_to_smvgc* function of the MVGC toolbox. Granger causality values were calculated for all integer frequency values within the desired range for all directed pairs of brain regions in the dataset.

For calculating electome network using, the exponential of all Granger causality values was used, which gives a ratio of total power to 'unexplained' power. Since the original formulation involves logarithms, it hinders the addibility of the features, so the exponential is suitable for inclusion in the electome model. Specifically,

$$\exp\left(\mathrm{f}_{\mathrm{Y}\rightarrow X}(\lambda)\right) = \frac{|\mathrm{S}_{\mathrm{XX}}(\lambda)|}{\left|\mathrm{S}_{\mathrm{XX}}(\lambda) - \mathrm{H}_{\mathrm{XY}}(\lambda)\Sigma_{\mathrm{Y|X}}\mathrm{H}_{\mathrm{XY}}(\lambda)^{*}\right|}$$

where $f_{Y\rightarrow X}(\lambda)$ represents Granger causality at frequency $\lambda$ from region *Y* to region *X*, $S_{XX}(\lambda)$ represent the spectral power in region *X* at frequency $\lambda$, and $H_{XY}(\lambda)\Sigma_{Y|X}H_{XY}(\lambda)^{*}$ represents the component of that power that is predicted by region *Y*. We capped values for this ratio at 10 to prevent any non-physiological signal from dominating the electome factors we learned using dCSFA-NMF.

**Discriminative Cross-Spectral Factor Analysis – Nonnegative Matrix Factorization**

To apply our Supervised Cross-Spectral Factor Analysis – Nonnegative Matrix Factorization (CSFA-NMF) model, which fully described elsewhere (Talbot et al., 2020), we consider each window of data to be an independent stationary measurement. This implies that the relevant dynamics happens at the scale of windows, so the extracted electome scores are all that is needed for later analysis. In this work, we choose a 1 second window because this balanced fine-grained behavior with enough length of signal to estimate the relevant LFP features. Prior work has shown relative robustness to windows between .5s to 5s in similar methods (Ulrich et al., 2015), so we expect similar results for similar window lengths; however, 5s here would not be able to capture the short-term scale of behavior necessary for this analysis.

For each window of data, we have the generated features, consisting of spectral power features, coherence features, and exponential granger features, totaling $P$ distinct features per window. Using the subscript $n$ to denote window and state that there are $N$ total windows. We describe the preprocessed data as $X_n \in \mathbb{R}_{+}^{P}$ (the $P$-dimensional non-negative domain) and the observed behavioral label as $y_n \in \{0,1\}$, where the binary indicates a social or non-social behavioral label. To briefly described this model, we set up an objective function to learn the $K$ different electome factors,

$$\min_{W,d,\phi} \sum_{n=1}^{N} \|x_n - Wf(x_n;\phi)\|_2^2 + \lambda\|y_n - d^T f(x_n;\phi)\|_2^2,$$

where each electome is described by a column in $W \in \mathbb{R}_{+}^{P\times K}$ (e.g., $W = [w_1, \ldots, w_K]$), the electome factor scores are given by the multi-output function $f(x_n;\phi)\colon \mathbb{R}_{+}^{p} \rightarrow \mathbb{R}_{+}^{K}$, and the relationship between the electome factor scores and the behavioral labels is given by $d \in \mathbb{R}^{p}$. The relative importance of reconstructing the observed data and the importance of the predictive task were balanced by choosing the hyperparameter $\lambda$. This represents a novel method to fit an NMF model using supervised autoencoders and requires the user to choose a parametrization for $f(x_n;\phi)$. In our method, this is simply set to an affine function following by a non-linearity, $f(x_n;\phi) = softplus(Ax + b)$, where the parameters of the function are $\phi = \{A, b\}$ and the softplus means an element-wise operation of the operation $\mathrm{softplus}(a) = \log(1 + \exp(a))$, which maps a real number to the non-negative space. This function can vary in complexity to allow greater model complexity, but we found that this function was sufficient in practice. Because this objective function follows a supervised autoencoder structure, a common deep learning structure, we are able to implement this technique in Tensorflow (Abadi et al., 2016) using the ADAM algorithm for learning (Diederik and Ba, 2014).

A benefit of using this structure for learning is that performing statistical inference from new data is fast and straightforward. In factor models, one typically has to set up an optimization algorithm to find the maximum a posteriori estimate. However, in our supervised CSFA-NMF framework, we can calculate the electome scores on new data simply by calling the function $f(x_n;\phi)$, allowing easy portability and facilitating future real time applications.

**Hyper-parameter Selection**

The proposed CSFA-NMF procedure requires us to choose several different settings in the algorithm, which was done with a cross-validation procedure where complete mice from the training set were left out. The hold-out mice, as described in the manuscript, were *not* used for hyperparameter selection and represent a true blind test set. Specifically, we must choose the number of electome factors $K$, the importance of the supervised task $\lambda$, the relative importance of the power features, coherence features, and exponential Granger features, and the parameterization of the mapping function $f(x_n; \phi)$.

We had dual goals in our analysis: reconstructing the original data well, which is to say that the learned Electomes actually describe the neural measurements well and predicting the behavioral task well. The reconstruction error was evaluated by the Mean Squared Error on the validation mice, and the performance on the behavioral task was evaluated by the mean Area Under the Curve (mean AUC) on the validation mice. Greater emphasis was placed on the behavioral task, so for each candidate number of electome networks $K$, we used the cross-validation procedure was used to choose the settings that maximized the mean AUC. After that, an elbow analysis was used to choose the number of electome networks $K$, which is to mean we choose the $K$ after which minimal gains in explaining the observed data was observed.

**Two-chamber social Interaction test**

Social preference was measured using a two-chamber assay in which animals explored a novel object or a novel mouse. The apparatus was a rectangular arena (61cm × 42.5cm × 22cm) constructed from clear plexiglass with a clear plexiglass wall dividing the arena into two equal chambers with an opening in the middle allowing free access between both chambers. The floor of the arena was constructed using a one-way mirror that allowed for video recording from beneath in order to avoid obstruction from electrophysiological recording equipment. Plastic, circular holding cages (8.3cm diameter and 12cm tall) were centered in each of the two chambers and were used to house either a novel object or sex- and age-matched C3H target mouse. The arena was evenly lit with indirect white light (~125 lux). Test mice were handled and habituated to the social preference chambers and empty holding cages for a least three days prior to testing. Subsequently, mice underwent ten separate social preference test sessions, with at least one day off in between sessions, in which the test mice were allowed to freely explore the arena for ten minutes; the holding cages contained either a novel object or novel C3H target mouse. The side of the chamber holding the object/mouse was determined pseudorandomly, such that the object/mouse would not be placed in the same chamber on more than two consecutive sessions in order to prevent side biases and to distinguish target-specific effects from location-specific effects. Plastic toys and glass objects were used as novel objects with the object being between 3-5cm in all directions. Video data was tracked using Bonsai Visual Reactive Programming software and the time spent in the proximity (5.8cm) of either holding cage was used to determine social preference scores.

The social preference for each session was defined as:

$$\frac{\overline{InteractionTime_S} - \overline{InteractionTime_O}}{\overline{InteractionTime_S} + \overline{InteractionTime_O}}$$

where $\text{Interactions}_S$ is the total time spent proximal to the other mouse, and $\text{InteractionTime}_O$ is the total time spent proximal to the object.

**Free Object/Social Interaction Test**

The Free Object/Social Interaction Test (FOSIT) allowed for free exploration of either novel objects or novel sex-matched conspecific mice during a single session. Plastic and glass objects were used similarly to social preference testing. The test was run in a clear arena (35cm × 31cm) lit using indirect white light (125 lux). The test mouse was placed into the arena that contained either a novel object or a target mouse and allowed free exploration (i.e., the objects/mice were not kept in holding cages as in the social interaction test) for five minutes. Following this five-min trial, the test mouse was placed into a new, identical arena that contained either a novel object or novel mouse for another five minutes. The order of object/mouse trials was determined pseudorandomly, such that the test mouse would not see a novel object or mouse for more than two consecutive sessions in order to prevent habituation to the stimulus type. Additionally, in order to control for the location of the target mouse, the novel object was pseudorandomly placed in one of four quadrants of the area such that each subsequent object placement was in a different quadrant from the previous object trial and that each quadrant was used at least once. A total of ten trials were run so that each test mouse was able to interact with five novel objects and five novel mice over the course of the ~50-min session. The amount of time interacting with the objects and target mice was hand-scored by experienced raters. For social trials, interactions were distinguished based on physical engagement (i.e., reciprocal interaction, test mouse investigating the target mouse, and target mouse investigating the test mouse).

**Spike-*Electome Factor* activity correlation**

Data acquired during the fifth session of the two-chamber social interaction test were used for this analysis. Cellular firing activity was averaged within one-sec non-overlapping windows for the ten-min recording period. The social firing preference of each cell was defined as:

$$\frac{\overline{FR_S} - \overline{FR_O}}{\overline{FR_S} + \overline{FR_O}}$$

where $FR_S$ is the neuronal firing rates observed during social interactions and $FR_O$ is the neuronal firing rates observed during interactions with an object. A rank-sum test of all one-sec observations was used to determine if a cell signaled social vs. object interactions. We used a spearman rank correlation to quantify the relationship between cellular firing during the ten-minute sessions and *Electome Factor* activity. We performed 1000 permutations for which Electome activity time bins were randomly shuffled within the social and object conditions. We then calculated the spearman rank correlation for each permutation. A cell was deemed to be positively correlated with the *Electome* network if it exhibited a spearman Rho above the 97.5% of the permutated distribution, and negatively correlated if it was below the 2.5%.

**Sucrose consumption**

Neural responses to sucrose or water delivery were measured in a rectangular chamber (30cm × 19cm × 28cm) constructed from black plastic Legos. Two nose poke holes, spaced 6.5cm apart along one of the long walls, detected nose pokes via IR beam breakage and delivered 10μL of fluid from a 27-gauge syringe situated within the hole; a five-sec timeout followed each fluid delivery in which subsequent nose pokes were not rewarded. Mice implanted with electrodes were habituated to the fluid drinking apparatus for two days prior to electrophysiological recordings. During habituation, singly housed mice had *ad libitum* access to food and used the nose poke holes for access to water. Subsequently, electrophysiological recordings were collected during two fluid-drinking sessions: one for sucrose and one for water. Specifically, four hours into the dark-cycle, mice were recorded for 1.5 hours while poking for administration of a 2% sucrose solution from both poke holes. Immediately following sucrose administration, the sucrose was switched out and water was delivered through the poke holes for an additional two hours. Timestamps for each nose poke were synchronized and stored alongside electrophysiological data.

**Elevated plus maze test**

The elevated plus maze (EPM) has been previously described. Briefly, the EPM consists of four cross-shaped arms (30.5cm length× 30.5cm width, at 91.4cm height from floor) and a 5cm×5cm central region. Two 'closed' arms are surrounded on three sides by walls of 16.5cm height and the other two 'open' arms are surrounded by a short piece of tape approximately 1 mm in height. Mice were habituated to the behavioral room for two hours, 24 hours before testing. Following a one-hr habituation period on the test day mice were placed in the center region of the elevated plus maze facing a closed arm. Neural recordings were obtained for ten minutes, and the location of the mice was captured using video recordings. All EPM testing was performed at 50 lux.

**Optogenetic manipulation of the prelimbic → nucleus accumbens circuit element**

Ten-week only mice were anesthetized with 1.% isoflurane and placed in a stereotaxic device. A 33-gauge Hamilton syringe was used to infuse 0.5 µl of AAV2-CaMKIIa-Chr2-EYFP (Murugan et al., 2017)vector at a rate of 0.1 µl/min, bilaterally, into prelimbic cortex (1.8mm AP, ±0.5mm ML, 2.5mm DV from the skull) and the syringe was left in place for ten minutes following the injection. In order to mitigate pain and inflammation related to the procedure, all animals received carprofen (5 mg/kg, s.c.) injections once prior to surgery and then once every 24 hours for three days following viral injections.

Eight weeks after viral surgeries, mice were anesthetized again, and recording electrodes were implanted as described above. A fiberoptic cannula was built into the nucleus accumbens bundle (Hultman et al., 2016; Kumar et al., 2013). The tip of the 100µm diameter fiberoptic (Doric Lenses) was situated 400µm above the tip of the recording microwires in the core of the accumbens. *In vivo* recordings and stimulations were conducted after 5–6 weeks of recovery. We delivered light stimulation at 1–7mW bilaterally (473nm wavelength, LaserGlow, LRS-0473-GFM-00100-05; 589nm wavelength, LaserGlow, LRS-0589-GFF-00100-05), and the laser output was verified using a Power meter (Thorlabs, PM100D). We first stimulated a group of mice (n=3) at 7mW to mirror a prior study(Murugan et al., 2017). All these animals exhibited seizures (observed behaviorally and confirmed using LFP recording). Repeat stimulation several days later at 1.5mW also induced seizures in these mice. We tested two additional stimulation naïve mice at 1.5mW, and one of these mice exhibited seizures as well. Thus, all experiments presented in the main manuscript were performed in mice with no prior stimulation. Mice were stimulated at 1mW bilaterally. Several mice showed ictal activity in the cortical channels at the onset of stimulation. Ictal activity was accompanied by immobility, backwards walking, and grooming in these mice. Ictal activity (restricted to the cortical channels) and behavioral responses subsided spontaneously usually within 5-10 seconds of stimulation onset, at which time mice became behaviorally activated and showed increase social investigation.

Recordings were performed in six blocks during a single session. In each block mice were exposed to a new object and a new mouse. Each block lasted 7.5 minutes, during which mice were stimulated with blue light for 2.5 minutes and yellow light for 2.5 minutes, with 1.25 minutes between each stimulation period. The order of blue vs. yellow light stimulation and social vs. object exposure was pseudorandomized for each mouse, such that animals never experienced the same color light stimulation first for all three social or object blocks. Additionally, the order of the social vs. object exposures were pseudorandomized for each block such that mice never experienced all three social or object exposures first.

Two mice exhibited global LFP responses to stimulation and were removed from further analysis. One mouse did not show any LFP responses to stimulation in nucleus accumbens and was thus removed from further analysis. For analysis, behavior was hand-scored from video recordings as described above by experienced raters blind to laser conditions (blue vs. yellow). We combined behavioral and neurophysiological measures for all periods in which the subject mouse was engaged with the other mouse (unilateral or bilateral). EN-Social activity observed during these social encounters was normalized to the activity observed during object encounters and compared across blue and yellow light stimulation trials. Periods in which the experimental partner mouse was unilateral engaged with the subject mouse were excluded.

### QUANTIFICATION AND STATISTICAL ANALYSIS

#### Electome Model Fitting

The statistical analyses for the Electome model were performed using Python 3.6 and Tensorflow version 1.09. We used machine learning to define a single relevant electome. The total number of Electomes was chosen to minimize the reconstruction loss with the minimal number of factors as defined previously. The reconstruction loss was weighted such that each mouse and each condition were weighted equally. The supervision loss weighting was determined as the amount of entropy contained in the binary variable of social vs object score. The supervision strength was started at a low value and gradually annealed to the final value.

While multiple Electomes networks were learned from the training data, only one electome network's activity was predictive of the social activity. Therefore, when applied to the test set, only this electome was evaluated statistically, so no multiple comparisons corrections were required (a major advantage of such a factor model formulation). The predictive ability for each mouse was quantified using the area under the curve using the network strengths.

#### Validation Testing

For validation testing, we projected LFP data recorded from new mice and/or new paradigms into our initial learned *Electome* network model. We then performed direct comparison across conditions (e.g., behavioral conditions, genotypes, etc.) using the median *Electome* network activity score for each condition per mouse. Activity scores were compared using non-parametric statistics, or parametric statistics a Box–Cox transformation was applied the raw data. To further enable evaluation of the robustness of our findings, the decoding strength (area under the curve of the receiver operating characteristic, which takes into account the activity scores for all of the transformed time windows) was also provided in the main text alongside the statistical results obtained through direct comparisons of the median activity scores. For optogenetic validation studies we normalized network activity observed social interactions to the network activity observed during object interactions for each stimulation type (blue vs. yellow). We utilized this strategy because the 10 Hz signal induced in cortex and striatum had the potential to diminish our detection of *EN-Social* (which reflected organized LFP patterns at frequencies including 10Hz). A longer-term solution will require the implementation of NMF-based statistical approaches that reconstruct *EN-Social* activity from data sets in which a subset of areas are excluded (see our prior model based on gaussian processes as an example)(Gallagher et al., 2017).

DATA AND SOFTWARE AVAILABILITY

The code base for dCSFA-NMF analysis can be found at https://github.com/carlson-lab/encodedSupervision.
